# Molecular Characterization of *Fischerella uthpalarensis*, the first subsection V cyanobiont from a tropical *Azolla* species containing dual nitrogenases

**DOI:** 10.1101/2023.03.07.531633

**Authors:** B.L.D Uthpala Pushpakumara, Dilantha Gunawardana

**Author notes:** Correspondence /.

## Abstract

There have been theories presented on *Azolla* cyanobionts, known for voracious nitrogen fixation, vertical transmission of cyanobiont and helping transform a greenhouse planet to an icehouse one ~49 Mya. One such theory encapsulates the existence of two cyanobionts, named Major and Minor. We show here the identity of a possible minor cyanobiont of *Azolla* named *Fischerella uthpalarensis*. A likely cyanobiont with straight or curved filaments that were truly branched was isolated. Seven gene fragments, namely 16s rDNA (Forward and Reverse), RNA Polymerase, ITS1 region (Forward and Reverse), *nifD* and GroEL genes were utilized to identify the isolated cyanobiont. The best match based on BLASTn search tool was found in the RNA Polymerase beta subunit (*rpoC*) gene fragment, that showed 99.54% identity with 55% coverage to *Fischerella muscicola*. Phylogenetic inferences with the *rpoC* genetic locus and the GroEL protein sequence suggest a likely *Fischerella* genus identity. Furthermore, *VnfDG* and *VnfN* fragments too were amplified using PCR and sequenced to demonstrate that this cyanobiont has alternate nitrogenase genes, on top of the molybdenum counterpart, providing an advantage in lifestyle. We encountered a higher level of genomic-level synonymous substitutions, which was not reflected in protein sequences, namely *VnfDG* and *VnfN* gene products, which may be due to codon heterogeneity. We also propose for *F. uthpalarensis* atypicality in codon usage due to the likely acquisition of the V-nitrogenase operon from a presumed recent horizontal gene transfer (HGT) event. The cyanobiont from this study showcases a higher preference for AT over GC at the *VnfDG* composite locus again hinting at a symbiotic lifestyle.

## 1. Introduction

*Azolla* is a genus of independent, floating, aquatic, pteridophyte water ferns belonging to the *Azollaceae* family. *Azolla pinnata* belongs to section *Rhizosperma. Azolla* fronds harbor a mutualistic symbiosis with a filamentous, heterocyst-forming, nitrogen-fixing cyanobacterium most popularly named *Nostoc azollae.* The cyanobacterium is found within a narrow region in the extracellular ovoid cavity, which is ensheathed by a mucilaginous network[1], [2] (Qiu and Yu, 2003; Pereira and Vasconcelos, 2014). The relationship between *Azolla* and cyanobacterium is believed to have originated millions of years ago as a permanent plant-cyanobacterium symbiosis. *Azolla* has also been studied for its use in agriculture as a nitrogen biofertilizer[3], [4] (Wagner, 1997; Pabby et al., 2003). The genome of *Azolla filiculoides* was sequenced in 2018; this fern species may be useful as a substitute for urea and as a solution to climate change[5] (Li et al., 2018).

The theory of an exclusive cyanobiont living in *Azolla* has now been overtaken by the “two strain” hypothesis, whereby major and minor contenders, both of the genus *Nostoc* or *Anabaena* result in a subsection IV cyanobacterium[2], [6], [7] (Zimmerman et al., 1989; Gebhardt and Nierzwicki-Bauer, 1991; Pereira and Vasconcelos, 2014). It is estimated that the cyanobionts only make up 1% of the biomass of Azolla fronds [8](Rai et al., 2000) Lectins produced by plants, enabling recognition by the cyanobiont at lectin interfaces, are heterogeneous in *Azolla* species[9] (McCowen et al., 1986) suggesting to us that there are distinctive, varied and specific cyanobionts for different *Azolla* species. This alludes to distinctive entry points for diverse cyanobionts and a gamut of host specificities [10](Papaefthimiou et al., 2008), of which all contenders are perhaps yet to be discovered.

A few studies have led to successful isolation of the secondary cyanobiont[6], [7], [11]–[13] (Ladha and Watanabe, 1982; Zimmerman et al., 1989; Gebhardt and Nierzwicki-Bauer, 1991; Rajaniemi et al., 2005; Sood et al., 2008). Separation of the cyanobiont from *Azolla* is important for its identification using molecular biology tools. The identity of the cyanobiont of *Azolla* spp still remains uncertain[2] (Pereira and Vasconcelos, 2014); aside from the major-minor hypothesis of two *Nostoc/Anabaena* strains, there may be another genus living symbiotically inside *Azolla*[2], [14], [15] (Caudales et al., 1992; Ekman et al., 2008; Pereira and Vasconcelos, 2014). It is widely accepted that the major cyanobiont is a species of *Nostoc azollae;* however, the identity of a secondary cyanobiont has been unclear until this study.

Cyanobacteria and cyanobionts have been reported to possess 3 types of nitrogenases, the most common being the molybdenum-dependent nitrogenase, the second most common being the vanadium-reliant nitrogenase and the rarest, the iron-based nitrogenase [16](Hu et al., 2012). So far, vanadium nitrogenase dependent microorganisms have been discovered in wood chips, soils, and mangrove sediments, which show that they have the potential to arise from diverse environments[17] (McRose *et al.,* 2017). Outside of the genus *Azotobacter,* there are also, a handful of other soil bacteria that fix nitrogen with the help of vanadium, namely the genera, *Methylocysis, Phaeospirillum, Rhodomicrobium, Rhodopseudomonas, Methanosarcina* and *Tolumonas.* It has been calculated that alternative (V) nitrogenases contribute 14-21% of diversity of nitrogenases, and a 24% contribution, to nitrogen fixation, which extends their importance to rice growing soils[17] (McRose *et al.,* 2017). In cyanobacterial symbioses, namely the genera *Peltigera* (lichen), *Anthoceros* and *Blasia,* the latter two being bryophytes, biology appears to favor the presence of a vanadium-nitrogenase on top of the molybdenum counterpart [18](Nelson, 2019).

The fact that there is higher levels of vanadium in the soil appears to have little repercussions on the favorable expression of the molybdenum counterpart. In the genus *Azotobacter*, vanadium is transported inside using siderophores such as V-azotochelin only when the cells are conditioned to low molybdenum concentrations, which suggests that the internalization of vanadium to be coupled to the preferential activity of the V-dependent nitrogenase. Such internalization is dependent on “vanadophores”, a subset of siderophores that are able to transport vanadium internally[19] (Rehder, 2008). It has been demonstrated that the vanadium-dependent nitrogenases use ABC transporters for the internalization of soluble and bioavailable vanadate, in microorganisms such as *Anabaena variabilis*[20] (Pratte et al., 2006).

In this study, we report evidence for the first subsection V cyanobiont in an endosymbiotic relationship with a plant (likely to be *Azolla microphylla).* The sequence of experiments performed in this study include: the isolation of the cyanobiont, detailing the technical replicates and negative controls used to authenticate the isolation procedure; morphological investigation of the cyanobiont using optical microscopy; and the sequencing of five separate loci, followed by BLASTn analyses of fragments against the NCBI non-redundant nucleotide database. We have also reported in a recent paper that this cyanobacterium is likely to possess an “alternate” vanadium nitrogenase in addition to the molybdenum counterpart[21] (Pushpakumara and Gunawardana, 2018), to which we present genomic evidence.

## 2. Materials and Methods

### 2.1 Culture of the *Azolla* Cyanobiont in a Free-living State

The isolation and cultivation of the cyanobiont of *Azolla* plants in a free-living state was performed according to a previous protocol [21](Pushpakumara and Gunawardana, 2018). The protocol is explained in brief in the following paragraph. The isolation procedure was carried out inside a standard laminar flow. The working surface and walls of the laminar flow cabinet were cleaned using 70% ethanol before starting the isolation procedure.

Fifteen grams of fresh fern tissue was weighed and washed in running tap water for 10 minutes. The ferns were then surface sterilized in 10% Clorox for two minutes, followed by a one-minute rinse in sterile 0.01 M HCl. The ferns were then rinsed for one minute in sterile distilled water twice. Following the disinfection, ferns were homogenized in 50 mL BG11_0_/8, filtered, then the filtrate was centrifuged at 500 × g for 5 minutes. The resulting cell pellet was washed in 2 mL BG11_0_/8 and 500 μL was transferred to 10 mL BG11_0_/8 in a McCartney bottle. The medium was incubated under natural conditions, exposing an approximate 12-hour day and 12-hour night cycle for two weeks with intermittent shaking by hand. After two weeks incubation, 5 mL of culture was transferred to 10 mL BG11_0_/8 and incubated under the same conditions for another two weeks. After two weeks, 10 mL of culture medium was introduced to an increased volume (20 mL) of BG11_0_/8 in a conical flask and incubated on a gyrotory shaker at 150 rpm close to a window to receive sunlight during daytime. After 1 week of incubation, the culture was exposed to full strength N-free BG11o, where 3 mL of full-strength N-free BG11o was introduced in 5-day intervals until there was a significant biomass growth.

### 2.2 DNA Extraction

DNA extraction was performed using a modified protocol [22](Wilson, 2001).

### 2.3 PCR amplification and sequencing

PCR primers and conditions for PCR reactions of this study are provided respectively in Table S1 and S2 in supplementary materials. Amplified PCR products were analyzed using 1.5% agarose gel electrophoresis. The size of the band was confirmed using a 1 Kb ladder (Promega). Sequencing was performed bidirectionally using the respective forward and reverse primers at Macrogen (Korea).

### 2.3 Sequence Alignments and Phylogenetic Analyses

Sequence alignments were performed using ClustalW Omega server for both DNA and protein sequences[23] (Sievers et al., 2011) using the default parameters. A maximum likelihood tree for 22 gene sequences was constructed using MEGA software, (version X) [24][Sohpal et al., 2010], comprising downloaded sequences from subsections IV and V of cyanobacterial taxonomy. Relationships between adjacent nodes were based on bootstrap support from 500 pseudo-replicates. The phylogenetic tree was constructed following the multiple sequence alignment of the sequences using the default matrix (ClustalW) in MEGA X software.

### 2.4 Disulfide Bond Prediction

The selected sequence was searched against the DiANNA server (http://clavius.bc.edu/~clotelab/DiANNA/) for the identification of likely disulfide bond pairs[25] (Ferre and Clote, 2005).

## 3. Results

### 3.1: Deciphering Identity of an Isolated Cyanobiont: Isolation, Identification using Molecular Data, and Morphology

The Azolla-major cyanobiont hypothesis describes the partnership based on mutualism as long-standing, dating back hundreds of millions of years[26] (Hall and Swanson, 1968). The major cyanobiont has been given the generic identities of Nostoc, Anabaena and Trichormus, and is putatively undergoing genomic erosion and pseudogenization[2], [27] (Ran et al., 2010; Pereira and Vasconcelos, 2014). Scientists believe that it has either lost autonomy or will during epochs to come. Only a few gene families have been retained by the eroded genome, while the functionality of genes responsible for glycolysis and nutrient uptake have been lost [27](Ran et al., 2010). The genomic erosion of the major cyanobiont has made its independent culture difficult, if not impossible. The degree of gene loss has led to a poor outlook for culturing the major cyanobiont as a monoculture. It is noteworthy that the major cyanobiont is vertically transmitted, unlike other plant endosymbionts, which are horizontally transmitted[28] (Zheng et al., 2009). Therefore, there is no infection/colonization of the host; instead, the cyanobiont is inherited inter-generationally.

We initially investigated the cultivation of a cyanobiont, past literature has called “Minor”, in comparison to the uncultivable “Major” cyanobiont. For facility, we call this isolate “Minor” due to solely its cultivable status, and to demarcate it away from the major partner, which we know as Nostoc azollae. We are in no way suggesting exclusively the candidacy of the minor cyanobiont here, only suggesting an easy identifier, to a cultivable microorganism. It is noteworthy that many studies of the Anabaena/Nostoc/Trichormus-Azolla symbioses have attempted to separate the two perceived organisms and culture the cultivable cyanobacterium in a free-living state. Although isolation methods have been successful, there has been disagreement over the appropriate surface sterilization procedures used for Azolla, which requires closer investigation[2] (Pereira and Vasconcelos, 2014). In the present study, a surface sterilization procedure was used on 15 g (fresh weight) of fern tissue, which was initially washed under running tap water for 10 min. Ferns were then surface-sterilized in 10% Clorox for two minutes and rinsed for 1 min in sterile 0.01 M HCl. The ferns were then rinsed twice in sterile distilled water for 1 min. We also performed three control experiments (technical replicates), namely 5 to 6 fronds, which were transferred to a McCartney bottle containing 10 mL of one-eighth-strength N-free BG11o medium (N-free BG11_0_/8) under sterile conditions. The bottle was closed, and hand-shaken intermittently for 15 minutes. Fronds were then carefully removed using aseptic technique. The bottle containing 10 mL N-free BG11_0_/8, was then incubated under natural conditions (approximately 12 hours day and 12 hours night cycle) for two weeks and hand shaken 2-3 times per day. A green-colored growth was observed after 2 weeks in the surface sterilized and crushed fronds, while there was no growth in negative controls[21] (Pushpakumara and Gunawardana, 2018). This demonstrates that the isolate we describe later as a subsection V cyanobiont is not a transient epiphytic colonizer, but is a permanent endosymbiont. To date, only a single occurrence of a Fischerella species has been reported in the leaf compartment, which was found in a phyllosphere in the rain forests of Costa Rica[29] (Finsinger et al., 2008).

A key question is why no subsection V species have previously been reported from the leaves of Azolla species. The level of care devoted to the isolation process in this study and strength of the disinfection procedure ensure that the present findings are authentic. Furthermore, negative controls failed to yield a detectable culture. A recent study showed the use of a two-step approach to collect other minor bacterial inhabitants from the leaf cavity of Azolla (1% bleach for 40 seconds and four rinses of water) [30](Dijkhuizen et al., 2018). The present study details a more comprehensive approach consisting of four steps: water, Clorox, HCl, then water. The success in isolating cyanobiont from 3 technical replicates provided strong evidence that the cyanobiont is a permanent candidate that resides abundantly in the leaf cavity.

We grew the isolated cyanobiont and extracted DNA from the culture. From the genomic DNA template, the loci 16s rDNA (1432–1439 bp), RNA polymerase C (861 bp), ITS region (variable) nifD (338 bp) and GroEL (1485 bp) gene fragments were amplified for identification and taxonomic purposes. 16s rDNA is the gold standard for eubacterial taxonomy and we assessed this locus using both NCBI nucleotide database. It is noteworthy to mention that cyanobionts are the only nitrogen fixers found within Azolla, eliminating the possibility of other cyanobacteria aiding the formation of fixed nitrogen stocks[30]–[32] (Lindblad et al., 1991; Braun-Howland et al., 1998; Dijkhizen et al., 2018).

According to bacterial taxonomic rules, >97% identity shared between 16s rDNA sequences is sufficient for the classification of a bacterium at the species level [33](Tindall et al., 2010). Further, >95% is the upper boundary for estimating a genus; the lower boundary is >90%, although the lower boundary is not frequently used for taxonomic purposes [34], [35](Janda and Abbott 2007; Sabat et al., 2017). Initially, the nifD DNA fragment and cell morphology studies were used to determine the species of the cultured cyanobiont. The nifD contig derived from the forward and the reverse sequences, shared >92% sequence identity with Fischerella sp. UTEX 1903 (Table S3). The occurrence of polymorphisms varies between 16s rDNA and nifD genes; it is estimated that the polymorphisms in nifD genes are 6-fold greater than 16s rDNA genes, which makes them more variable and prone to mutate in related sequences[36] (Qian et al., 2003). A 396bp rpoB sequence has been shown before to be able to discriminate between Mycobacterial species which are 91.8% similar in sequence, suggesting that the usage of the nifD gene (338bp) to discriminate between members of IV and V cyanobacteria, is just as strong in phylogenetic resolve[37] (Devulder et al., 2005).

The 16s rDNA sequence obtained from post-amplification sequencing, pointed to a Fischerella muscicola relation, with the query sequences in forward (B23S primer) and reverse (pA primer), aligning with 61%-65% coverage to a similar locus in Fischerella muscicola when BLASTn was used for the determination of close relatives. The level of identities for 16s rDNA were 95%-97% which was sufficient for genus level identification but not for the determination of species level match. In fact, the >800 bp partial fragment of rpoC presented us with a 99.54% identity (at 55% coverage) with the same free-living microorganism, again suggesting that the likely cyanobiont was closest in taxonomy to the genus Fischerella. However, the sequencing data from all five loci (seven reads in total) were insufficient for the identification of the cyanobiont at the species level. The genome of Fischerella muscicola, isolated from a paddy field in India in 1951, has been sequenced and is likely to present the closest identity to the cyanobiont we have isolated.

RNA polymerase genes (rpoB and rpoC) are implicated in superior growth rates in E.coli, and consequently can be subjected to growth maximizing mutations. While the closest match of the rpoC contig assembled in this study, was Fischerella muscicola, there was a considerable portion of the sequence that was not covered by the sequenced 861 bp fragment. This segment (1-349 bp) did not align with any sequence in the NCBI nucleotide database. It has been demonstrated that large-scale genome rearrangement and variation in genes and gene content, are key factors in cyanobiont genomes containing vanadium-dependent nitrogenases. When a Maximum Likelihood phylogenetic tree was built using rpoC gene sequences from 22 cyanobacteria, rooted to the subsection IV, Nostoc sp. LP11, we could clearly establish the evolutionary relationship between the cyanobiont of this study and Fischerella muscicola as they formed a monophyletic clade, with 100% bootstrap support (Figure 1). There was a clear bifurcated separation of subsection IV and subsection V in the resulting phylogenetic tree.

**Figure 1:**
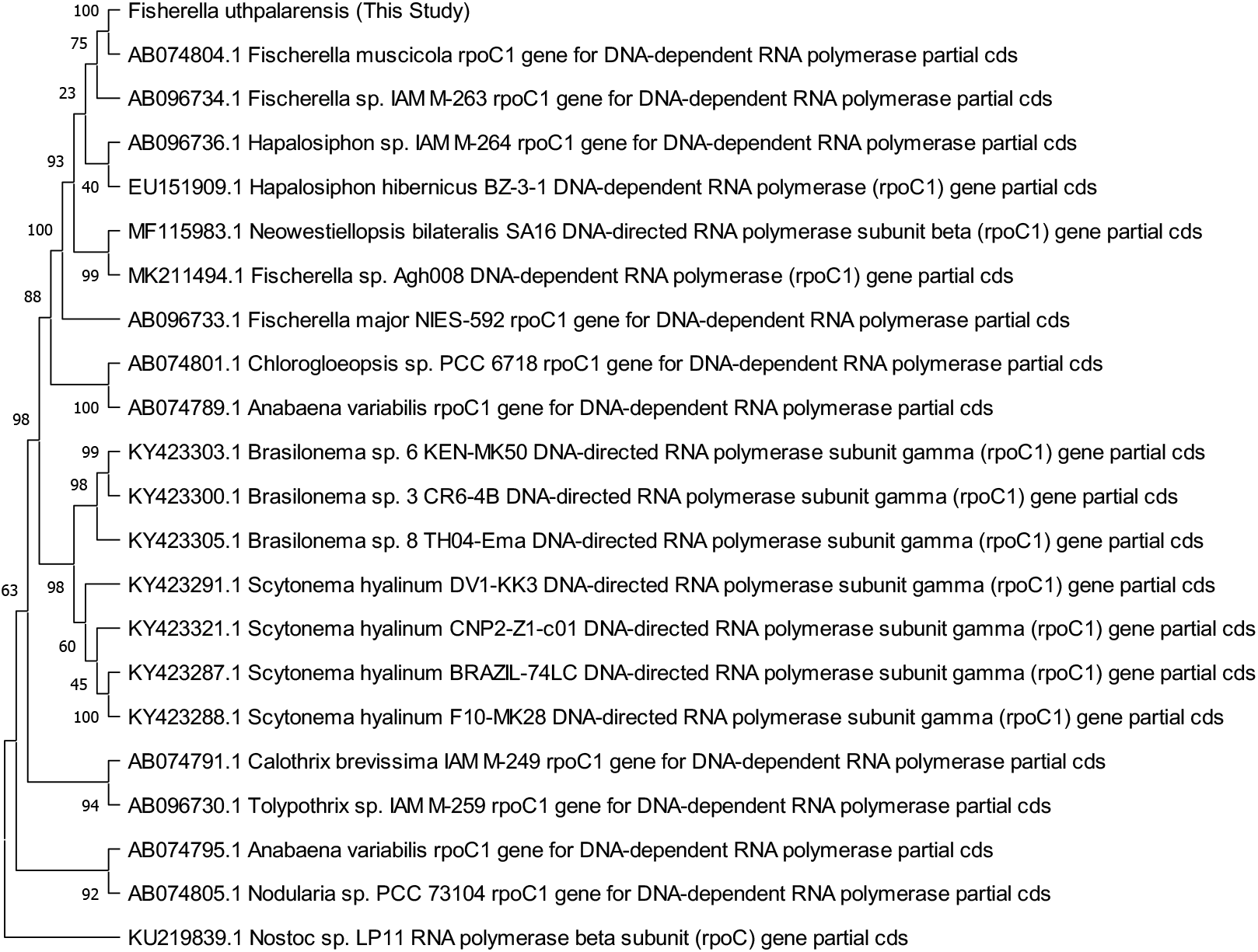
The Maximum Likelihood (ML) tree of the rpoC gene fragment of the isolate (*Fischerella uthpalarensis*) against 22 homologs from the NCBI nucleotide database constructed following sampling with 500 bootstrap pseudo-replications.

The internally transcribed spacer region is found between 16s and 23s rRNA sequences, which can be easily amplified using primers that bind to the conserved flanking sequences. The ITS region can be strongly variable which transforms such regions to be good candidates for phylogenetic inferences at higher taxonomic levels. The region sequenced by the ITSCYA236F primer encompasses a region that is 240 bp is length within the locus AF105134 of *Fischerella muscicola,* representing a region of tRNA-Ile and tRNA-Ala genes (incomplete sequences), internal transcribed spacer (partial sequence) and 23s ribosomal RNA (partial sequence). The region sequenced by the ITSCYA236F reverse primer is representative of a similar fragment; tRNA-Ile and tRNA-Ala genes (incomplete sequences), internal transcribed spacer (partial sequence), and 23s ribosomal RNA (partial sequence) spanning from 9 to 250, of the AF105134 (460 bp total sequence) fragment. Cyanobacteria possess three kinds of 16s-23s ITS regions, the most prevalent being the ones that contain the two tRNA genes, that has been detected in *Nostoc, Anabaena* and many other Nostocales cyanobacteria, which is perhaps synonymous with the findings of this study. We too assessed phylogeny from GroEL proteins since they are included/listed as loci/products capable for phylogenetic inferences (Figure 2). In the study by Xie et al., 2019, there were several genes listed that the authors assessed as candidates for gene or protein based phylogeny inferences, namely groEL, rplB, rpoB, recA, tuf, aroE, ddl, dnaE, fusA, ftsZ, glnA, gyrB, glpF, gltX, gyrB, gp d, gdh, hemN, ileS, lepA, leuS, ldhL, mutS, mutL, metRS, nrdD, pepV, pgm, polA, recG, recP, xpt, yqil, tkt, and tpi, where they say mention GroEL gene as a unitary candidate capable of rapid mutational change and consequently a boon for phylogeny inferences [38](Xie at al., 2019).

**Figure 2:**
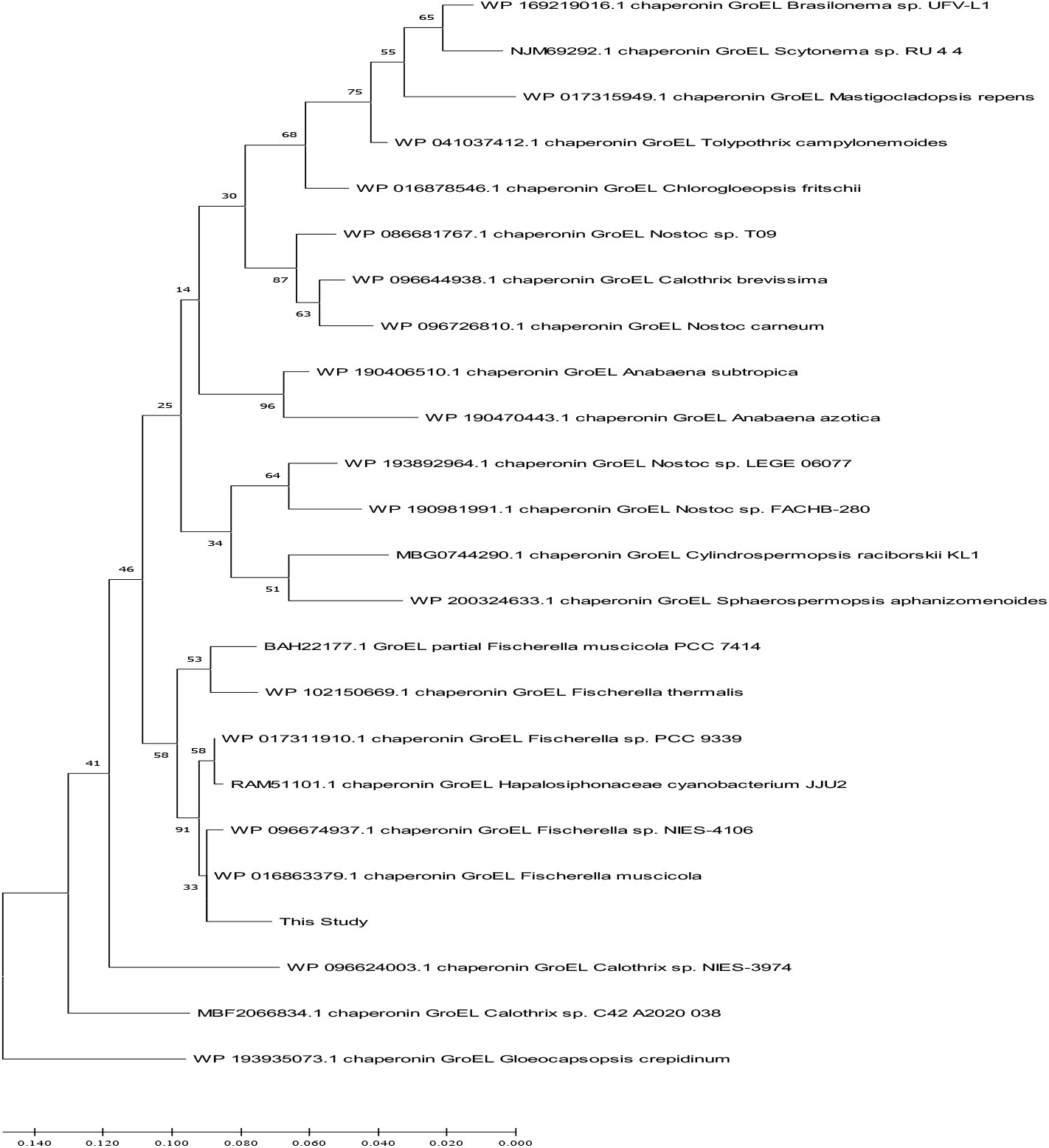
The Maximum Likelihood (ML) tree of GroEL proteins of the isolate (*Fischerella uthpalarensis*) against 23 homologs from the NCBI nucleotide database constructed following sampling with 1000 bootstrap pseudo-replications. Gloeocapsopsis crepidinum was assigned as outgroup.

When the phylogeny was inferred using the GroEL translated protein with 23 other GroEL sequences from cyanobacteria, using Gloeocapsopsis crepidinum as the outgroup, the cyanobiont sequence focused in this study formed a monophyletic cluster with four Fischerella sequences and one Hapalosiphonaceae cyanobacterium JJU2 sequence, which had 91% bootstrap support for their overall monophyly, with the most immediate node of the cyanobiont belonging to Fischerella muscicola though with only 33% bootstrap support. Hapalosiphonaceae cyanobacterium JJU2 is a subsection V cyanobacterium isolated from the Philippines, which is a true-branching and heterocyst-forming bacterium, that is resistant to ambient cadmium[39] (Victoria et al., 2018). A separate clade consisted of falsely-branching cyanobacteria including Brasilonema sp. UFV-L1, Scytonema sp. RU_4_4 and Tolypothrix campylonemoides, clustered with true-branching Chlorogloeopsis fritschii and Mastigocladopsis repens, which showed again the polyphyly of stigonematales cyanobacteria, into two distant clusters[40] (Gugger and Hoffmann, 2004). The paddy-field inhabiting, vanadium nitrogenase equipped species Anabaena azotica and Anabaena subtropica, were found as a single clade with 96% bootstrap support of their phylogenetic positions as proximal neighbors. Overall, inner phylogeny was supported by high bootstrap values but this was not the case for outer branches.

Species of the genus Fischerella are known to have filamentous, heterocystous, barrel-shaped cells. Fischerella belongs to subsection V of cyanobacterial taxonomy, which is different in lineage from Anabaena or Nostoc, which belong to subsection IV. True branching patterns (either T, V, or Y branching anatomy, which are symbolic of subsection V cyanobacteria), were clearly observed in isolated cyanobiont’s microscopy images (Figure 3). Such cyanobacteria undergo transverse, oblique, and longitudinal cell divisions[40] (Gugger and Hoffman, 2004). It can also be seen that cell division occurs in many planes, as in the genus Fischerella, while forming V-shaped junctions with a “keystone or bridging cell” at the center that anchors three-way branching (Figure 3). On the other hand, other Fischerella species (e.g. F. muscicola) possess T-shaped junctions that branch at perpendicular angles[40] (Gugger and Hoffman, 2004). To summarize, sequencing and morphology results suggest that a novel subsection V cyanobiont in a plant symbiosis, the first of its kind, has been isolated in this study.

**Figure 3:**
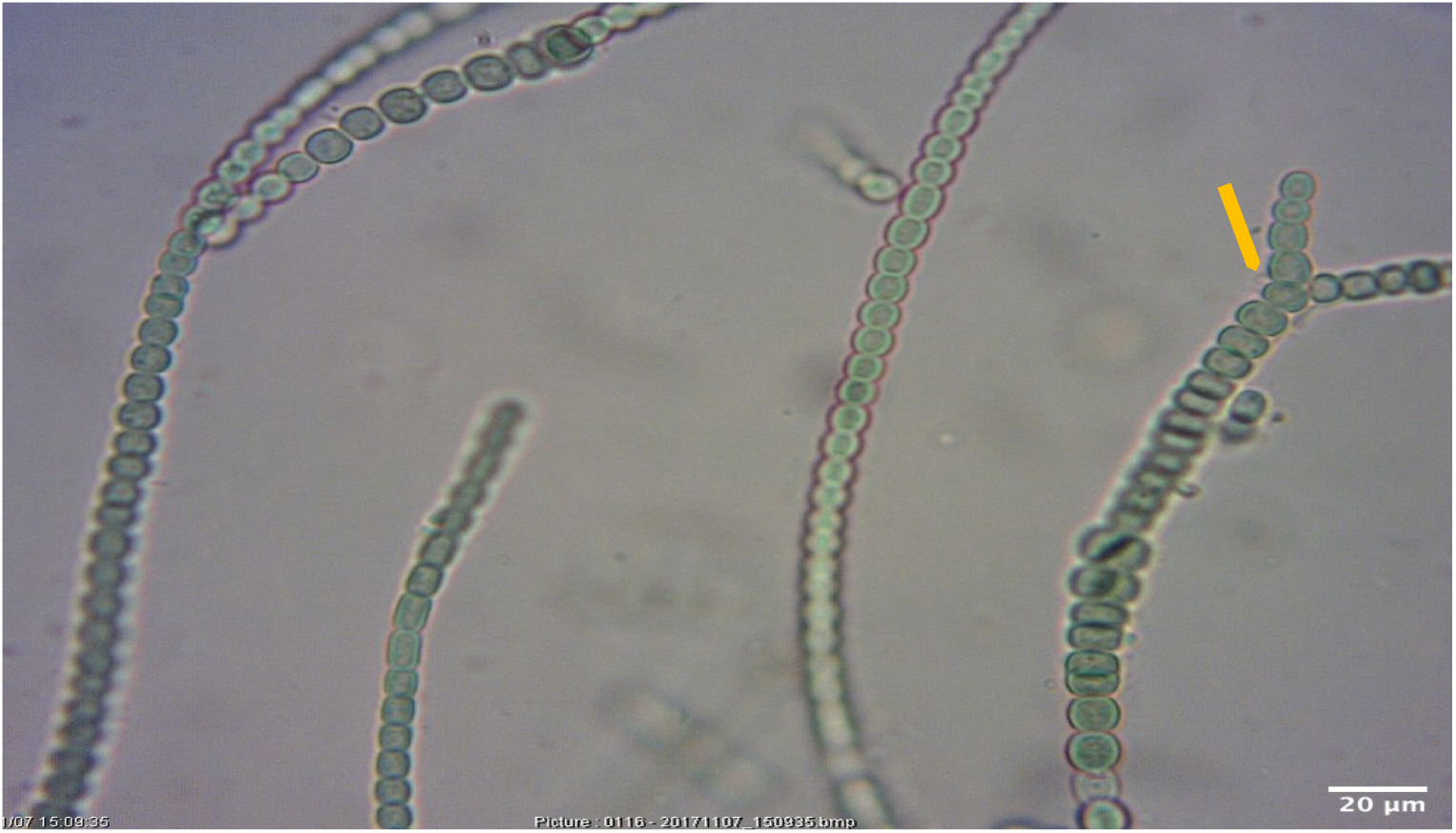
Branching patterns of the isolate (×400) visualized using optical microscopy. The already branched forms are indicated **by an orange arrow**. Branches originate from a “keystone” like central cell that attaches to three distinct cells from three separate branched filaments

### 3.2: Dual Nitrogenases of Fischerella uthpalarensis

We showed in two previous articles, that the minor cyanobiont inside a likely tropical *Azolla* microphylla, has a likely vanadium-dependent nitrogenase, on top of the molybdenum counterpart[21], [41] (Atugoda et al., 2018; Pushpakumara and Gunawardana, 2019). We show here conclusively using both PCR amplification and sequencing of two loci, that an alternate vanadium nitrogenase is encoded by the cyanobiont genome. The PCR products from vnfDG (A fusion single gene made of vnfD and vnfG) and VnfN were synonymous with the expected size of the amplicons, while sequencing showed that they encoded for parts of an alternate vanadium nitrogenase (Figure S1(A) and (B)). We too have amplified nifH and nifD genes from the cyanobiont’s genome (Figure S1 (C)) showcasing both nitrogenases are present.

It has been demonstrated that cyanobionts of Peltigera lichens and bryophyte genera, Blasia and Anthoceros, have cyanobionts with vanadium-dependent nitrogenase-encoding capacities in their genomes, and our study adds another cyanobiont to the growing list of cyanobionts proven to encode vanadium nitrogenases[18] (Nelson, 2019). Complementation of function by both host and infecting cyanobiont is crucial for the establishment of a symbiosis. Out of 4 Nostoc cyanobionts of bryophytes, three contain the genetic background to produce vanadium dependent nitrogenases, two of which have the vanadium nitrogenase genes in plasmid genomes pointing to horizontal gene transfer as means of genetic exchange for the transfer of a new function (Nelson et al., 2019). Vandium nitrogenases are catalytically-distinct and are produced by separate genetic regions of the respective cyanobiont genomes. In this study, we were able to amplify the vnfDG and vnfN genes from DNA isolated from *Fischerella uthpalarensis*.

When the vnfN gene fragment sequenced in this study, was searched using the BLASTn tool, the best alignment was found in Nostocales cyanobacterium HT-58-2 at a paltry 65% sequence coverage and 88.34% sequence identity, while the second-best alignment was found with Peltigera membranacea (LA31632_AccXBB013(IINH)) cyanobiont, at 65% coverage and 85.88% identity (Table S4). There were no candidates from the genus Fischerella in the list of homologous gene fragments when the DNA sequence was used for BLASTn searches. In contrast to the BLASTn results, when the vnfN fragment was translated in all six open reading frames (ORFs), the longest translated sequence gave us a protein product that had 95% sequence coverage and 99% sequence identity to the vnfN protein from *Fischerella muscicola* when searched using the BLASTp search tool (Table S4).

A similar story was shown for the bidirectionally sequenced amplicon produced by VnfDG2F and VnfDG4R primers, again showing 82.32% identity with 82% coverage to the nearest match, while when the longest ORF was translated, the translation product showed 98.48% sequence identity to Fischerella muscicola (Table S4). A higher deviation in both the vnfN and vnfDG amplicons against the nearest neighbor at the DNA level was not reflected in the sequence deviation in relation to “protein sequence” queries, which yielded a near perfect match (98-99% identity); i.e. synonymous substitutions are commonplace in sequence. The explanation to this observation can entail several possibilities from genome instability, differences in generation time (how fast recombination occurs giving rise to mutations), codon usage heterogeneity, DNA-repair systems and host-supplied proteins that may have impacts on the types of mutations[42] (Lopez et al., 2019).

The rice plant (Figure 4) in particular, is known to be surrounded by a few vanadium dependent N-fixing microorganisms. Both, Anabaena azotica FACHB-118 and Anabaena sp. CH1, associated with rice paddies contain an alternate nitrogenase [43]. Furthermore, bacteria of the genus Azotobacter are widely cultured from paddy fields and such bacteria employ V-dependent nitrogenases for nitrogen fixation. Vanadium can be found as four oxidation states that are relatively stable in aqueous environments, namely V^2+^, V^3+^, VO^2+^, and VO^2+^ (the oxidation states 2, 3, 4, and 5, respectively).

**Figure 4:**
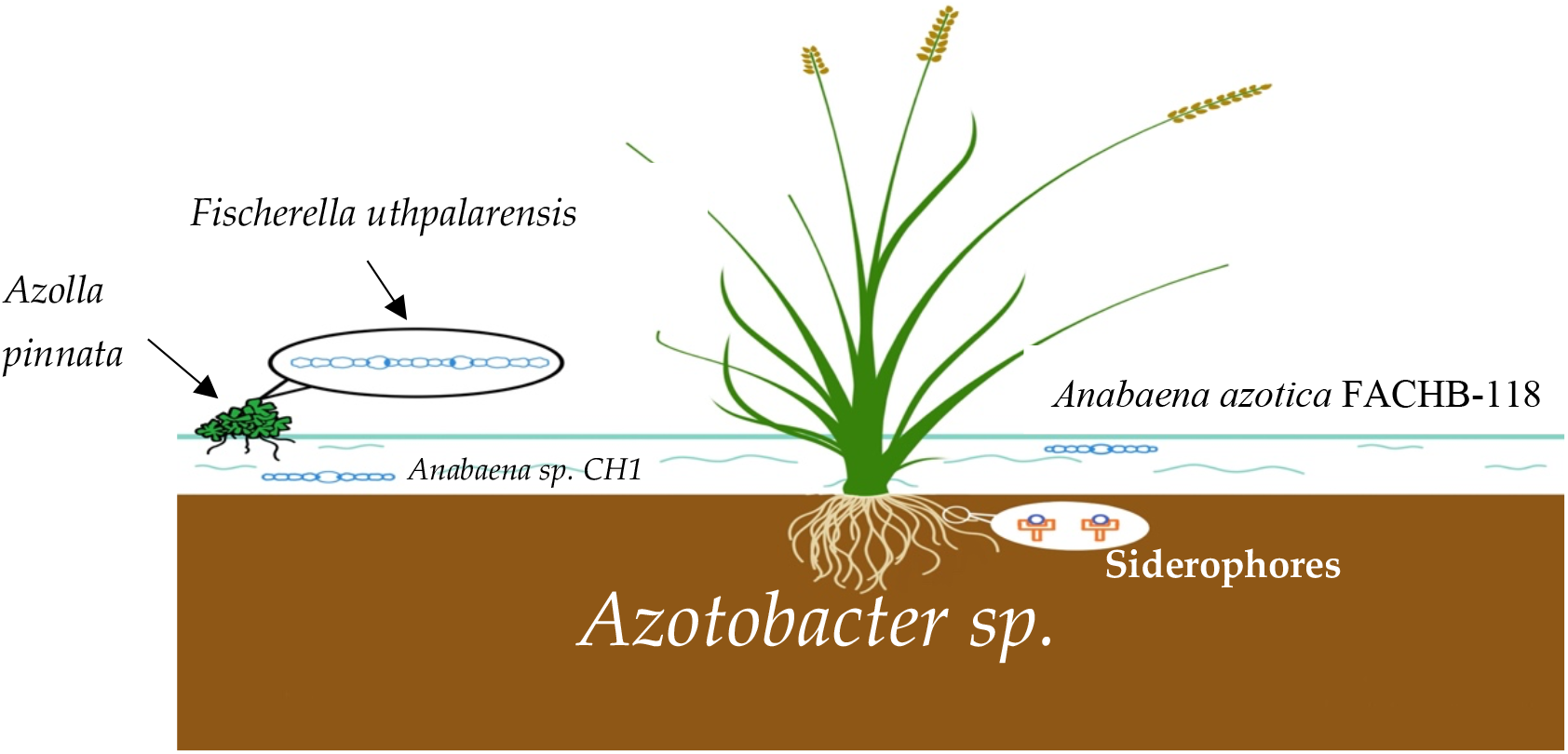
The illustrated irrigated ecosystem of a rice plant showcasing the vanadium-dependent microorganisms

Vanadium is approximately 1-100 fold higher in the soil compared to molybdenum. Molybdenum on average, is found at 1-10 ppm in Asian paddy soils while measured as 20-30 ppm in a subset of Asian soils. The mean molybdenum content in tropical paddy soils is within the bracket of 2.3 – 3.3 ppm out of which the lowest value was found in Sri Lanka, West Malaysia and Cambodia[44] (Domingo and Kyuma, 1983). In contrast, a separate study measures vanadium content as between 3 to 500 ppm[45] (Swaine 1955, Bowen, 1979). The analysis of vanadium in tropical Asian soils, gives a mean value of 166 ppm that is within the above ranges[44] (Domingo and Kyuma, 1983). Whether such skewed numbers of the ratio between vanadium and molybdenum contribute to the opulence of V-nitrogenase armed microorganisms in paddy cultivated areas, is a future empirical crusade. Chemically, Vanadium is redox-sensitive and therefore prone to be transformed to a reduced redox potential in flooded environments such as paddy fields. A higher solubility of vanadium is likely to occur under most flooding conditions due to vanadate(V)-sorbing iron complexes attenuated in dissolution capacities and due to vanadate(V) being reduced to vanadyl (IV) that forms complexes with organic matter in irrigated rice fields [46](Gustafsson, 2019). Therefore, the availability of vanadium increases with irrigation/flooding which may be the reason for the richness of V-dependent microorganisms in paddy fields.

### 3.3: Molecular clues as to Horizontal Gene Transfer and Symbiosis

We draw upon “alien/atypical” codon usage to gather evidence of horizontal gene transfer events. It is hypothesized that recent horizontally-acquired genes have “alien/atypical” codon usage and undergo an amelioration period from donor to recipient lifestyles following the transfer event[42] (Lopez et al., 2019), which we infer here for the VnfDG and VnfN genes. For example, E.coli evolved from a linage of Salmonella ~100 Mya, and during that window, E.coli has been the recipient of 234 lateral transfer events[47] (Ermolaeva, 2001). Furthermore, highly expressed genes such as nitrogenases are dependent on the availability and abundance of the correct tRNA species for translation of mRNA to a fully-fledged protein [48](Wang et al., 2011). Highly expressed genes such as VnfDG are under higher natural selection for highly efficient codons.

We looked at the functionally-key cysteine (6 in number in the partial fragment) within the protein sequence of the VnfDG gene product. In analyzing codon usage heterogeneity (Table S5), we found evidence as to the higher utilization of the TGT codon for cysteine (over the TGC counterpart), which was atypical in relation to the common usage of the two codon types (Figure S2(A) and (B); Table S5). Such a phenomenon was not observed in free-living Anabaena variabilis full length VnfDG gene where TGC>TGT in coding for cysteines (data not shown). In fact, three TGT codons (out of a total four) encoding cysteines in F. uthpalarensis VnfDG partial sequence are found as either TGC (2) and CGT (1) in the model organism for alternate nitrogenases, A. variabilis (Figure S2 (B)).

Cysteines are crucial from a functional perspective and were not available at the origin of life. Cysteines are equipped with thiol/sulfhydryl groups, have a propensity to form disulfide bonds, are found as highly-conserved residues in protein sequences, forms clusters in close proximity, possess high metal binding affinities, while have a duality of opinion in relation to its hydrophobic/non-hydrophobic nature[49] (Poole 2015). Our observations propose that widespread DNA level synonymous substitutions can be attributed to codon heterogeneity stemming from different codon usage strategies in Fischerella uthpalarensis. Two cysteines in the VnfDG gene product of F. uthpalarensis, the authors propose here to be stemming from important mutations (figure S2 (B)). When the DIANNA web server was used to show function of the above Cys-98, the result revealed a case for a half-cystine (Cys-98), which are known to form disulfide bonds with downstream or upstream residues (Table S5).

The authors suggest the newly acquired nature of the alternate nitrogenase-encoding operon could mean that the codon usage preferences are yet to be reassigned within the new host, and consequently can be designated as atypical or alien. On average cyanobacteria have been recorded to have 36-78 tRNA genes per genome, although this number is 42 in obligate symbiotic Trichormus azollae, 58 in free-living Fischerella muscicola CCMME 5323, 70 in facultative symbiotic Nostoc cycadae WK-1 and 67 in facultative symbiotic V-nitrogenase-possessing Nostoc sp. ‘Peltigera membranacaea’ 210A (NCBI Nucleotide Portal). Fast growing bacteria have been shown to contain more tRNAs (median of 61) compared to slow-growing counterparts (median of 44) (Wang et al., 2011). There is too an example from the extreme end of symbioses – AT rich plastid genomes – where the psbA gene does not follow the host rules for codon usage, instead following an ancestral model of codon usage[50] (Morton and Levin, 1997). What is remarkable of the chloroplast psbA gene is that it is subjected to a high degree of synonymous substitutions compared all other chloroplast encoded genes – as is the case here for VnfDG and VnfN – but shows a remnant inventory of preferred codons, for the recruitment of the correct amino acids [50](Morton and Levin, 1997). In fact, in the AT-rich Euphorbiaceae chloroplast genomes, TGT is classed as a high frequency codon, which again strengthens our arguments on horizontal gene transfer events [51](Wang et al., 2020).

In addition, silent sites (synonymous substitutions) are suggested to originate to facilitate alterations in gene expression, and consequently, the basis for selection of synonymous base modifications are thought to be a measure for gene expression leading to stronger codon biases in highly-expressed genes[52] (Kudla et al., 2006). The stronger suite of synonymous substitutions in VnfDG (82%) and VnfN loci (88%) from this study suggests that they may be tinkered for better gene expression although there is no footprint on the functional VnfDG and VnfN gene products as effector molecules. There are higher levels of GC% in sites of synonymous substitutions, than adjacent regions, that may have repercussions on gene expression of highly expressed genes such as VnfDG[52] (Kudla et al., 2006). In summary, the enrichment of silent sites in VnfDG and VnfN may be candidates to relax/optimize gene expression.

Symbiotic islands in rhizobia (such as the loci of the nifHDK operon) have low GC% compared to non-symbiotic counterparts, where the third positions of codons, in particular, are subjected to conversion to be poor in GC content[53] (Okubo et al., 2016). While the free-living A. variabilis has 43.1% and 45.25% GC content in VnfN and VnfDG genes, this value is reduced in our isolate where the corresponding values are, VnfN (42.5%) and VnfDG (41.9%) (Table S7). The GC content too has an impact on codon heterogeneity and usage, especially when the wobble position is converted from G or C to A or T, as shown for TGT codons at the expense of TGC codons (Table S5). Trichormus azollae, an obligate symbiont, has a GC content of 38.3%. The obligate cyanobiont of the unicellular algae Rhopalodia gibba has a GC content of 40.8% for the region spanning the nif operon[54] (Kniep et al., 2008), while bacterial symbionts such as Buchnera sp. have reduced genomes with AT biases. There is also evidence as to the higher levels of GC% in nitrogen fixing genomes against those incapable of fixing nitrogen[55] (Hildebrand et al., 2010) which again reinforces our claims for the nitrogenase ORFs of Fischerella uthpalarensis, as relative paupers of GC content, and consequently be likely AT-rich candidates for an incumbent mutualistic symbiosis.

We propose here that Fischerella uthpalarensis provides a possible “symbiotic” system to study due to availability of the genome of the likely parent strain, Fischerella muscicola, and due to the cultivability of the microorganism from fronds of Azolla plants

## Supplementary Materials

**Figure S1(A):** Amplified PCR product *(VnfN* gene) using VnfN2F and VnfN4R primers. **(B)** Amplified PCR product *(VnfDG* gene) using VnfDG2F and VnfDG4R primers. The blue arrows are bands of 600 bp and 1500 bp.

**FigureS1(C):** PCR products of *nifD* and *nifH* genes amplified using nifD_For & nifD_Rev and PolFor & PolR respectively. Lane 1: DNA marker, Lane 2 and 3: *nifD* (338 bp) and *nifH* (360 bp) amplicons obtained from the DNA of the isolated free living minor cyanobiont, *Fischerella uthpalarensis*. Lane 4 and 5: *nifD* (338 bp) and *nifH* (360 bp) amplicons obtained from the pooled DNA of the *Azolla*-cyanobiont symbioses (whole plant DNA) respectively.

**Figure S2 (A):** Sequence alignment of a partial sequence of *VnfDG* locus of free-living *Anabaena (Trichormus) variabilis* and *Fischerella uthpalarensis.* The codons encoding for cysteines (as shown in Figure 5(A)) are highlighted (yellow/green) while all changed codons are shown in blue squares where there is an AT bias in *F. uthpalarensis.* 3 out of 6 codons (50%) in *F. uthpalarensis*, have replaced a “C” with a “T”, two such codons being changed at the wobble (third) position.

**Figure S2(B) Top:** The region containing the cysteine codons encoding the signature protein motif, Cys (98)-X-X-Cys (101). “Study” is the *VnfDG* locus from *F. uthpalarensis,* while the remaining sequences are the “closest matches” using BLASTn search tool.

**Figure S2(B) Bottom:** A similar alignment for the region spanning the 6^th^ cysteine encoded by TGT in the the *VnfDG* locus from *F. uthpalarensis.*

**Table S1:** Details of Primers used in this study.

**Table S2:** Details of PCR cycling conditions for each primer pair used in this study.

**Table S3**: The locus, primers and the BLAST data obtained using BLASTn search tool.

**Table S4:** BLAST searches of sequenced DNA of selective sequenced genes encoding vanadium dependent protein products and their translated counterparts.

**Table S5:** Prediction of half-cystine, free cysteines and ligand-bound cysteines in the partial *VnfDG* gene product of *F. uthpalarensis.*

**Table S6:** The codon usage for cysteine and lysine globally, in freshwater cyanobacteria and in *VnfDG* locus of our isolate. The global percentages were obtained from the following webpage https://www.genscript.com/tools/codon-frequency-table)

**Table S7:** The GC% content of *VnfDG* regions in three cyanobionts, *Anabaena variabilis* and the cyanobiont under scrutiny here in this study.

## Author Contributions

This study was performed by Mrs. B.L.D Uthpala Pushpakumara as an undergraduate research project under the supervision of Dr. Dilantha Gunawardana at the Department of Botany, University of Sri Jayewardenepura. Dilantha pursued the story of the cyanobiont using molecular biology while Uthpala Pushpakumara commenced her PhD in an area of marine symbioses. Uthpala and Dilantha, respectively a PhD student and an alumnus of University of Melbourne, were responsible for the conceptualization. Dilantha wrote the manuscript

## Data Availability Statement

The datasets generated during and/or analysed during the current study are available from the corresponding author on reasonable request.

## Acknowledgments

This study was an undergraduate research project at the Department of Botany, University of Sri Jayewardenepura. The authors thank Genetech, Sri Lanka for technical support.

