## Supplementary Materials for "Molecular Characterization of *Fischerella uthpalarensis*, the first subsection V cyanobiont from a tropical *Azolla* species containing dual nitrogenases"

**Table S1: Details of Primers used in this study.**

| Primer Name | Sequence (5’ → 3’) | Locus | Amplicon Size | Study |
|---|---|---|---|---|
| nifD_F | CGGTTACTGGTCTTGGTCTGGTC– | *NifD* | 338 bp | Glass et al., (2010) |
| nifD_R | GCGTCGTTAGCGATGTGGTGTC | *NifD* | 338 bp | Glass et al., (2010) |
| B23A | CTTCGCCTCTGTGTGCCTAGGT | 16s rDNA | 1432-1439 bp | Edwards et al., (1989) |
| pA | AGAGTTTGATCCTGGCTCAG | 16s rDNA | 1432-1439 bp | Lepere et al., (2000) |
| ITSCYA236F | CTGGTTCRAGTCCAGGAT | ITS region | Variable | Valerio et al., (2008) |
| ITSCYA225R | TGCAGTTKTCAAGGTTCT | ITS region | Variable | Valerio et al., (2008) |
| ROPC145F | ACTCTGAARCCAGAAATGGA | rpoC | 861 bp | Valerio et al., (2008) |
| RPOC1006R | TGCTTACCTTCAATAATGTC | rpoC | 861 bp | Valerio et al., (2008) |
| vnfDG1F | TATTAAAGTGCGACGAAAC | VnfDG | 1041 bp | Hodkinson et al., (2014) |
| VnfDG6R | CCATCATCAATATAGAT | VnfDG | 1041 bp | Hodkinson et al., (2014) |
| VnfN2F | AAAGATGTCAGTATTGT | VnfN | 1080 bp | Hodkinson et al., (2014) |

| | | | | |
|---|---|---|---|---|
| VnfN4R | GCCATGTATTTTTCCCA | VnfN | 1080 bp | Hodkinson et al., (2014) |
| VnfDG2F | ATCAAAGAAAARGGSGAAGA | VnfDG | ~900-1000 bp | Hodkinson et al., (2014) |
| VndGD4R | ATACAGACTTTTTTGCC | VnfDG | ~900-1000 bp | Hodkinson et al., (2014) |
| GroELfor | ATGGCAAAGCGCATTATCTACAACG | GroEL | ~1495 bp | (This Study) |
| GroELRev | ATCAA CGATACCAGC AGC | GroEL | ~1495 bp | (This Study) |

**Table S2: Details of PCR cycling conditions for each primer pair used in this study.**

| **Primer Pair** | **Conditions** |
|---|---|
| pA and B23S | 94 –10min<br>**94-2min**<br>**50-45sec**<br>**72- 2min**<br>72-15min |
| ITSCYA236F and ITSCYA225R | 94 - 5min<br>**94-45sec**<br>**46-30sec**<br>**72- 45sec**<br>72-5min |
| ROPC145F and ROPC145F | 94 - 5min<br>**94-1min**<br>**44-30sec**<br>**72- 1min**<br>72-5min |
| NifD_F and NifD_R | Glass et al., 2010 |
| vnfDG1F and VnfDG6R | Hodkinson et al., 2014 |
| VnfN2F and VnfN4R | Hodkinson et al., 2014 |
| VnfDG2F and VnfDG4R | Hodkinson et al., 2014 |
| GroELFor and GroELRev | Outsourced. |

**Table S3**: The locus, primers and the BLAST data obtained using BLASTn search tool.

| **Fragment** | **Primer used For Sequencing** | **Closest Match (Species Name)** | **Query Coverage** | **Levels of Identity to closest match** |
|---|---|---|---|---|
| nifD (Contig) | NifD_For and NifD_Rev | *Fischerella* sp. UTEX 1903 | 98% | 92% |
| ITS region (Forward) | ITSCYA236F | *Fischerella muscicola* | 10% | 95.42% |
| ITS region (Reverse) | ITSCYA225R | *Fischerella muscicola* | 23% | 97.52% |
| RNA Polymerase C (Contig) | RPOC145F and RPOC1006R | *Fischerella muscicola* | 55% | 99.64% |
| 16S ribosomal RNA gene and 16S-23S ribosomal RNA intergenic spacer (Forward) | pA | *Westiellopsis* sp. DW5 II | 66% | 94.58% |

| | | | | |
|---|---|---|---|---|
| ITS region/Partial ribosomal RNA operon (Reverse) | B23S | *Fischerella muscicola* SAG 2027 | 61% | 96.59% |
| GroEL | GroEL-For And GroEL-Rev | *Fischerella sp.* NIES-4106 | 98% | 96.08% |

**Table S4:** BLAST searches of sequenced DNA of selective sequenced genes encoding vanadium dependent protein products and their translated counterparts.

| **Sequence** | **DNA level BLASTn Query** | | | **Protein level BLASTp Query** | |
|---|---|---|---|---|---|
| | **Coverage** | **Identity** | | **Coverage** | **Identity** |
| **VnfN** | 65% | 88.34% | | 95% | 99% |
| **VnfDG** | 82% | 82.36% | | 100% | 98.48% |

**Table S5:** Prediction of half-cystine, free cysteines and ligand-bound cysteines in the partial *VnfDG* gene product of *F. uthpalarensis.*

| Sequence **STUDY** Length **264** residues | | | | |
|---|---|---|---|---|
| Cysteines in this sequence: **6** | | | | |
| Cysteine Class prediction | | | | |
| **Cysteine** | **Half-cystine** | **Free cysteine** | **Ligand-bound** | **Ligand** |
| 8 | 0.243109 | 0.594115 | 0.162776 | - |
| 71 | 0.110079 | 0.799226 | 0.090694 | - |
| 98 | 0.435852 | 0.377152 | 0.186996 | - |
| 101 | 0.299121 | 0.467293 | 0.233586 | - |
| 190 | 0.141743 | 0.779899 | 0.078357 | - |
| 220 | 0.100271 | 0.764817 | 0.134912 | - |

**Table S6:** The codon usage for cysteine and lysine globally, in freshwater cyanobacteria and in *VnfDG* locus of our isolate. The global percentages were obtained from the following webpage (https://www.genscript.com/tools/codon-frequency-table)

| Amino Acid | Codon | Number /Percentage In VnfDG (This study) | Global Percentage | In Freshwater Cyanobacteria (Phytolase genes) |
|---|---|---|---|---|
| Cysteine | TGT | 4 (0.66) | 0.46 | 0.36 |
| Cysteine | TGC | 2 (0.34) | 0.54 | 0.64 |

**Table S7:** The GC% content of *VnfDG* regions in three cyanobionts, *Anabaena variabilis* and the cyanobiont under scrutiny here in this study.

| Species | VnfDG GC% |
|---|---|
| This Study | 41.9% |
| *Nostoc* sp. *'Peltigera dolichorhiza'* | 46.3% |
| *Nostoc* sp. *'Peltigera membranacea'* | 44.5% |
| *Nostoc* sp. *'Peltigera malacea'* | 42.2% |
| *Anabaena variabilis* | 45.3% |

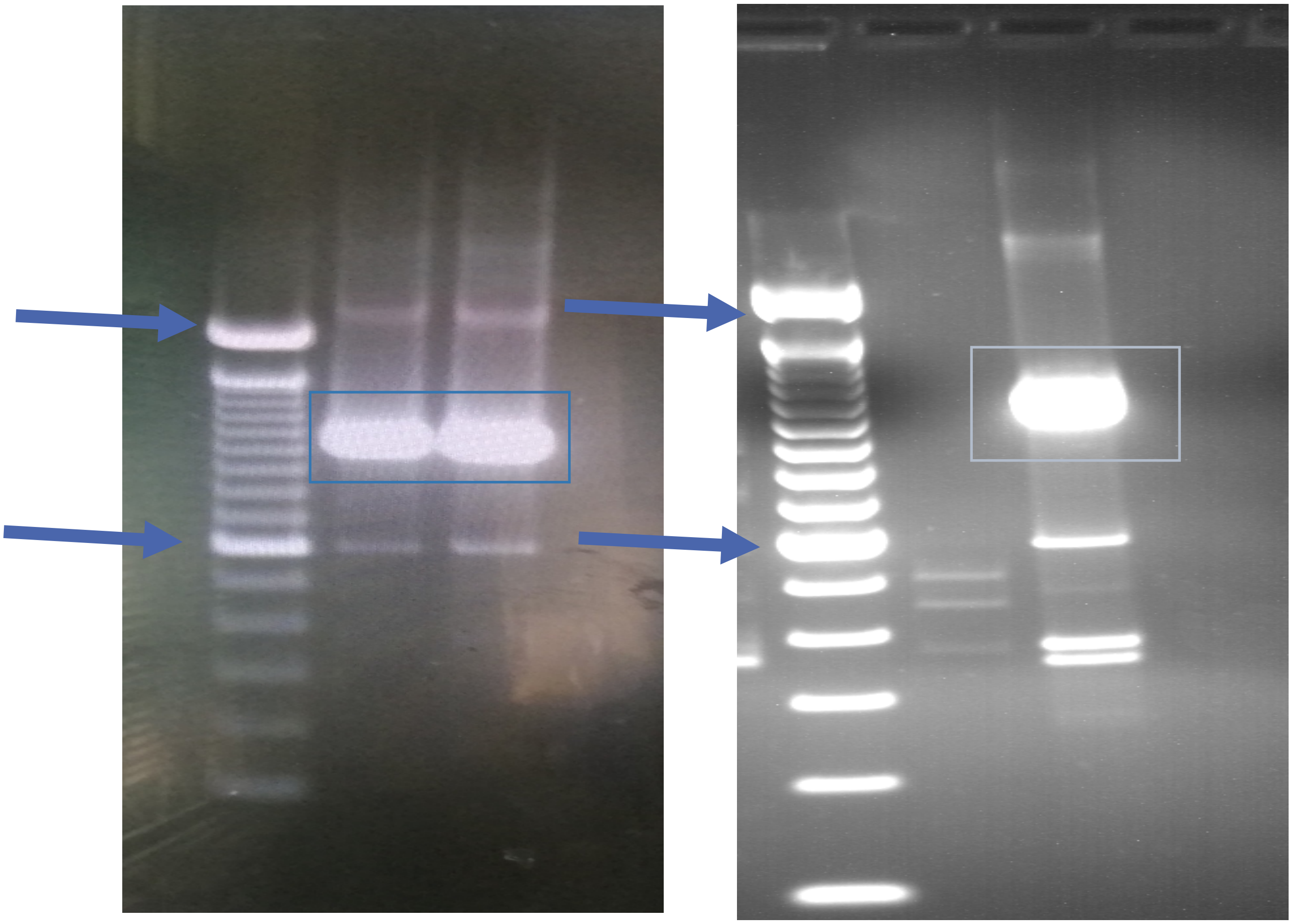

**Figure S1(A):** Amplified PCR product (*VnfN* gene) using VnfN2F and VnfN4R primers. **(B)** Amplified PCR product (*VnfDG* gene) using VnfDG2F and VnfDG4R primers. The blue arrows are bands of 600 bp and 1500 bp.

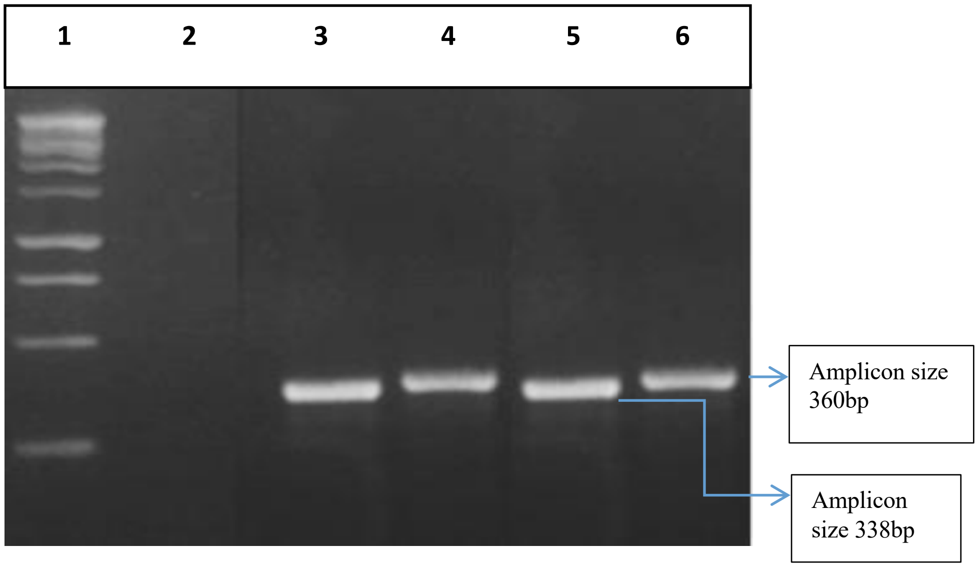


**FigureS1(C):** PCR products of *nifD* and *nifH* genes amplified using nifD_For & nifD_Rev and PolFor & PolR respectively. Lane 1: DNA marker, Lane 2 and 3: *nifD* (338 bp) and *nifH* (360 bp) amplicons obtained from the DNA of the isolated free living minor cyanobiont, *Fischerella uthpalarensis*. Lane 4 and 5: *nifD* (338 bp) and *nifH* (360 bp) amplicons obtained from the pooled DNA of the *Azolla*-cyanobiont symbioses (whole plant DNA) respectively.

```
Anabaena         1 atgcc------actaaaactattaaagtgc-ga-cgaaacaatcc----c   38
                   |..||      ||.|||||.|||   |.|| || ||.|||||.||    .
Fischerella     36 AATCCTAAAATACCAAAACGATT---GGGCAGATCGTAACAAGCCGTATA   82

Anabaena        39 agaaagggagaagcacgtatacatc-------------------------   63
                   |.|.|.||.|..|.||||..||..|
Fischerella     83 AAATAAGGCGGCGAACGTGCACCACCCGCCTCTTTTTTTTTTTTTTTTTT  132

Anabaena        64 aaagaaaaaggcgaagacacaacccaatttctgcccctatccaacatcga  113
                   |||.||||||  |||||||||||.||.|||||.||.||.||||||||||
Fischerella    133 AAAAAAAAAG--GAAGACACAACTCAGTTTCTACCTCTTTCCAACATCGA  180

Anabaena       114 aaccatcccaggttcactatcagaacgtggatgtagctactgcggagcca  163
                   |||.||.||.||||||||.|||||.||.||||||||||||||.||.||||
Fischerella    181 AACAATACCTGGTTCACTTTCAGAGCGGGGATGTAGCTACTGTGGTGCCA  230

Anabaena       164 aactggtaattggtggcgtactcaaagacaccatccaaatgattcacggc  213
                   ||||.||.||||||||.||.|||||||||||.||.||||||||||||.|||
Fischerella    231 AACTTGTGATTGGTGGTGTTCTCAAAGACACTATTCAAATGATTCATGGC  280

Anabaena       214 cccataggttgcgcctacgacacctggcacaccaaacgttatcctagcga  263
                   |||||.||||||.|||||||.||.||||||||||||||.||||||...||
Fischerella    281 CCCATCGGTTGCTCCTACGATACTTGGCACACCAAACGCTATCCTTCTGA  330

Anabaena       264 caacggtaactttcaactaaaatacgtctggtcatcagacatgaaagaat  313
                   ||||||.|||||.|||||.||.||.|..||||||...||||||||||.|
Fischerella    331 CAACGGCAACTTCCAACTCAAGTATGCTTGGTCAAGCGACATGAAAGAGT  380

Anabaena       314 cccacgtcgtcttcggtggagaaaaacaactaggcaaatccatccgagaa  363
                   |.||..|||||||||||||.|||||.|||||....|||||.||.|..|||
Fischerella    381 CGCATATCGTCTTCGGTGGCGAAAAGCAACTCAAAAAATCGATGCTGGAA  430

Anabaena       364 gccttcaaagaatttcccgacatcaaacggatgatagtctacaccacctg  413
                   |||||...||||||||||||.|||||.|||||||||||.||||||||||
Fischerella    431 GCCTTTGCAGAATTTCCCGAAATCAAGCGGATGATAGTTTACACCACCTG  480

Anabaena       414 cgccacagccttgattggtgacgacatcaaagccgtagtcaaaagcgcac  463
                   |||.||.||..|.|||||.||.||.|||||||||||.||.||||..|...
Fischerella    481 CGCTACTGCACTTATTGGAGATGATATCAAAGCCGTTGTTAAAAATGTCG  530

Anabaena       464 agcaagaactcggtgacgtagacatcttctgcgtagaatgtcccggattt  513
                   |..||.||||.|||||.|||||.|||||  ||.||.||.|||||.||.|||
Fischerella    531 AAAAACAACTTGGTGATGTAGATATCTTCTGTGTTGAGTGTCCTGGTTTT  580

Anabaena       514 gcaggtgtgagccaatccaaaggacaccacgtcctcaacatcgcttggat  563
                   ||||||||.|||||||||||||.||||||||.|||||.||.|..|||.|
Fischerella    581 GCAGGTGTCAGCCAATCCAAAGGCCACCACGTTCTCAATATTGGCTGGGT  630

Anabaena       564 taacgaaaaagtcggcacattagagccagaaatcacctcaccctacacca  613
                   .||.||.||.||.||.|||.|.|||||.|||||||||...||.||.||.|
Fischerella    631 GAATGAGAAGGTGGGAACACTTGAGCCGGAAATCACCAGTCCATATACGA  680

Anabaena       614 tcaacgtcatcggagactacaacatccaaggtgacaccttcgtactagag  663
                   |.||.||||||||.|||||||||||.|||||.||.|||||.||.||||||
Fischerella    681 TGAATGTCATCGGCGACTACAACATTCAAGGCGATACCTTTGTGCTAGAG  730

Anabaena       664 aagtacatggagaaaatgggcattcaaatcatcgcccattttaccggaaa  713
                   |||||||||||.||||||||..|.||||||||||||||||||||.||.||
Fischerella    731 AAGTACATGGAAAAAATGGGTGTGCAAATCATCGCCCATTTTACGGGTAA  780

Anabaena       714 tggtacttatgactccttacgaggaatgcacagagcgcaactcaacgtta  763
                   .|||||.||.|||||.||.|||||.||||||||||||||..|.|||||||
Fischerella    781 CGGTACATACGACTCATTGCGAGGTATGCACAGAGCGCAGTTAAACGTTA  830
```

```
Anabaena         764 ccaactgtgcgcgttcagccggatacatcgctaacgaactcaaaaagaga   813
                     ||||||||||||||...||.||||||||.||.||.||..|.||.||||||
Fischerella      831 CCAACTGTGCGCGTAGTGCTGGATACATTGCCAATGAGTTGAAGAAGAGA   880

Anabaena         814 tatggtattcctcgtctagacgtagacacctggggttttgactattgcca   863
                     |||||||||||.|||||.||.|||||.||.|||||.|||||||||||..|
Fischerella      881 TATGGTATTCCCCGTCTGGATGTAGATACTTGGGGCTTTGACTATTGTAA   930

Anabaena         864 agaagcattacgcaaaatcggcgcattcttcgggattgaagacagagccg   913
                     .||||||||.||.||||||||.||.|||||.||.|||||.||||||||||
Fischerella      931 GGAAGCATTGCGGAAAATCGGTGCCTTCTTTGGTATTGAGGACAGAGCCG   980

Anabaena         914 aggctgtaattgccgaagaaatcgccaaataccaagagaaaatggattgg   963
                     |.|||.|.|||||.||||||.|.||||||||..||...||..|.|..|||
Fischerella      981 AAGCTTTGATTGCTGAAGAAGTTGCCAAATATGAATCAAAGCTAGCGTGG  1030

Anabaena         964 tacaaggaaagactctcaggcaaaaaagtctgtatttggacaggtggtcc  1013
                     ||.||.|||.|..|....|||||||||.||||...|...|
Fischerella     1031 TATAAAGAACGTTTAAAGGGCAAAAAAATCTGATATAAAA----------  1070

Anabaena        1014 gagactatggcactggacaaaagctctggaagatgacttaggaatgcagg  1063
                      |.|..|....|....|.|||..|||||.|        ||
Fischerella     1071 -AAAAAAAAAAAAAAAAAAAATCCTCTGTA--------TT----------  1101

Anabaena        1064 tagtatccatgtcttctaagttcggacaccaagaagactttgaaaaggtc  1113
                        |.|...|.|||||    |||   |.||.|               .||
Fischerella     1102 ---TTTTTTTTTCTTC----TTC---CTCCTA---------------ATC  1126

Anabaena        1114 attgccagaggtcaagaaggaactatctatatt   1146
                     .|                    .|.|||.||.|
Fischerella     1127 CT--------------------ATTTCTTTAAT   1139
```

**Figure S2 (A):** Sequence alignment of a partial sequence of *VnfDG* locus of free-living *Anabaena (Trichormus) variabilis* and *Fischerella uthpalarensis*. The codons encoding for cysteines (as shown in Figure 5(A)) are highlighted (yellow/green) while all changed codons are shown in blue squares where there is an AT bias in *F. uthpalarensis.* 3 out of 6 codons (50%) in *F. uthpalarensis*, have replaced a "C" with a "T", two such codons being changed at the wobble (third) position.

```
Study          CGAAAAACAACTTGGTGATGTAGATATCTTCTGTGTTGAGTGTCCTGGTTTTGCAGGTGT 320
KF662362.1     AGAAAAAGAACTTGGTGACGTGGATATCTTCACTGTTGAGTGTCCCGGTTTTGCTGGAGT 431
KF662361.1     CGAAAAAGAACTGGGTGATGTGGATATCTTCACCGTTGAGTGTCCCGGTTTTGCTGGTGT 388
MN562834.1     CGAAAAAGAACTKGGTGATGTCGATATCTTCACCGTTGAGTGTCCCGGTTTTGCTGGTGT 467
MN562840.1     CGAAAAAGAACTKGGTGATGTCGATATCTTCACCGTTGAGTGTCCCGGTTTTGCTGGTGT 467
MN562827.1     CGAAAAAGAACTKGGTGATGTCGATATCTTCACCGTTGAGTGTCCCGGTTTTGCTGGTGT 491
MN562829.1     CGAAAAAGAACTTGGTGATGTCGATATCTTCACCGTTGAGTGTCCCGGTTTTGCTGGTGT 491
MN562824.1     CGAAAAAGAACTKGGTGATGTCGATATCTTCACCGTTGAGTGTCCCGGTTTTGCTGGTGT 491
KF662359.1     CGAAAAAGAACTTGGTGATGTAGACATCTTCACCGTCGAGTGTCCGGGTTTTGCTGGGGT 1260
MN562835.1     CGAAAAAGAACTTGGTGATGTCGATATCTTYACCGTCGAGTGTCCCGGTTTTGCTGGGGT 488
MN562839.1     AGAAAAAGAACTTGGTGATGTCGATATCTTCACCGTTGAGTGTCCAGGTTTTGCTGGTGT 491
MN562833.1     AGAAAAAGAACTCGGTGATGTCGATATCTTCACTGTTGAGTGTCCCGGTTTTGCTGGGGT 491
MN562821.1     AGAAAAAGAACTTGGTGATGTCGATATCTTCACCGTTGAGTGTCCCGGTTTTGCTGGTGT 491
                ****** **** ***** ** ** *****     ** ******** ******** ** **
```

```
Study            TCCCCGTCTGGATGTAGATACTTGGGGCTTTGACTATTGTAAGGAAGCATTGCGGAAAAT 680
KF662362.1       TCCCCGGATTGATGTGGATACCTGGGGCTTTGACTATGCCAAAGAAGGGTTACGCAAAAT 791
KF662361.1       TCCCCGGATCGATGTCGATACTTGGGGCTTTGACTATGCTAAGGAAGGGTTACGCAAAAT 748
MN562834.1       TCCCCGGATCGATGTCGATACTTGGGGCTTTGACTATGCTAAGGAAGGGTTACGCAAAAT 827
MN562840.1       TCCCCGGATCGATGTCGATACTTGGGGCTTTGACTATGCTAAGGAAGGGTTACGCAAAAT 827
MN562827.1       TCCCCGGATCGATGTCGATACTTGGGGCTTTGACTATGCTAAGGAAGGGTTACGCAAAAT 851
MN562829.1       TCCCCGGATCGATGTCGATACTTGGGGCTTTGACTATGCTAAGGAAGGGTTACGCAAAAT 851
MN562824.1       TCCCCGGATCGATGTCGATACTTGGGGCTTTGACTATGCTAAGGAAGGGTTACGCAAAAT 851
KF662359.1       TCCCCGGATTGATGTGGATACTTGGGGCTTTGACTATGCCAAGGAAGGGTTACGCAAAAT 1620
MN562835.1       TCCCCGGATTGATGTGGATACCTGGGGCTTTGACTATGCCAAGGAAGGGTTACGTAAAAT 848
MN562839.1       TCCCCGGATCGATGTCGATACTTGGGGCTTTGACTATGCTAAGGAAGGGTTACGCAAAAT 851
MN562833.1       TCCCCGGATCGATGTGGATACTTGGGGCTTTGACTATGCTAAGGAAGGGTTACGCAAAAT 851
MN562821.1       TCCCCGGATTGATGTGGATACTTGGGGCTTTGACTATGCTAAGGAAGGGTTACGCAAAAT 851
                 ******  * ***** ***** ***************   ** ****  ** ** *****
```

**Figure S2(B) Top:** The region containing the cysteine codons encoding the signature protein motif, Cys (98)-X-X-Cys (101). “Study” is the *VnfDG* locus from *F. uthpalarensis*, while the remaining sequences are the “closest matches” using BLASTn search tool.

**Figure S2(B) Bottom:** A similar alignment for the region spanning the 6$^{th}$ cysteine encoded by TGT in the the *VnfDG* locus from *F. uthpalarensis.*
